# Study of growth, metabolism, and morphology of *Akkermansia muciniphila* with an *in vitro* advanced bionic intestinal reactor

**DOI:** 10.1101/2020.07.06.190843

**Authors:** Zhi-tao Li, Guo-ao Hu, Li Zhu, Zheng-long Sun, Yun-Jiang, Min-jie Gao, Xiao-bei Zhan

## Abstract

As a kind of potential probiotic, *Akkermansia muciniphila* abundance in human body is directly causally related to obesity, diabetes, inflammation and abnormal metabolism. In this study, *A. muciniphila* dynamic cultures using five different media were implemented in an *in vitro* bionic intestinal reactor for the first time instead of the traditional static culture using brain heart infusion broth (BHI) or BHI + porcine mucin (BPM). The biomass under dynamic culture using BPM reached 1.92 g/L, which improved 44.36% compared with the value under static culture using BPM. The biomass under dynamic culture using human mucin (HM) further increased to the highest level of 2.89 g/L. Under dynamic culture using porcine mucin (PM) and HM, the main metabolites were short-chain fatty acids (acetic acid and butyric acid), while using other media, a considerable amount of branched-chain fatty acids (isobutyric and isovaleric acids) were produced. Under dynamic culture Using HM, the cell diameters reached 999 nm, and the outer membrane protein concentration reached the highest level of 26.26 μg/mg. This study provided a preliminary theoretical basis for the development of *A. muciniphila* as the next generation probiotic.

## 1. Introduction

In recent years, the research and the development of *Akkermansia muciniphila* has attracted interest[1–5]. *A. muciniphila* is an elliptical, immobile, strictly anaerobic Gram-negative strain that can use mucin in the intestine as carbon, nitrogen, and energy sources for growth[6]. The main metabolite of *A. muciniphila* are short-chain fatty acids (SCFAs) and branched-chain fatty acids (BCFAs) [7–10]. *A. muciniphila* was considered as a potential probiotic because evidences proved that *A. muciniphila* had a causal relationship with obesity[11–14], diabetes[15–17], inflammation[18–19], autism[20–21], amyotrophic lateral sclerosis[22], premature aging[23], epilepsy[24], hypertension[25], cancer[26] and metabolic abnormalities[27]. Recent studies reported that butyric acid produced by intestinal microbes like *A. muciniphila* could promote β cells to secrete insulin and then regulate blood sugar[28]. In addition, *A. muciniphila* cells with thicker outer membrane protein (Amuc-1100) showed better effects in preventing obesity and related complications[14].

In terms of *in vitro* culture of *A. muciniphila*, the traditional way was to use brain heart infusion broth (BHI) or BHI + porcine mucin (BPM) in an anaerobic bottle and placed it in an anaerobic environment for static fermentation culture[29–31]. Porcine mucin (PM) was the key component of *A. muciniphila* media in most reports. By adding a certain amount of mucin to BHI to cultivate *A. muciniphila*, the nutritional requirements for its growth and metabolism could be met. For simulation of *A. muciniphila* physiological state in human body, using human mucin (HM) instead of PM for *in vitro* culture is one key issue of great significance. However, as far as we know, the *in vitro* culture of *A. muciniphila* using HM has not been reported yet.

For culture method, the currently used *A. muciniphila* static culture is easy to cause accumulation of metabolites, which in turn inhibits the growth of *A. muciniphila.* In addition, *A. muciniphila* cells are difficult to harvest because many of them adhere to bottle bottom and wall. The *in vitro* dynamic fermentation reactor is a recently developed tool for studying the intestinal flora, mainly through bionic technology to simulate the real environment in digestive system [32]. In this study, an advanced bionic intestinal reactor (ABIR) was built to simulate the peristalsis of the real human colon, the micro-ecological environment of intestinal and the absorption of flora metabolites. As the traditional static culture state is far from the real *A. muciniphila* growth environment in human body, using ABIR as the *A. muciniphila* culture equipment is another key issue of great significance for simulating *in vivo A. muciniphila* physiological environment.

In this study, in order to better understand the growth, metabolism and morphology of *A. muciniphila* in real human body, *in vitro* dynamic *A. muciniphila* fermentation was implemented using five different media including HM and BHI + human mucin (BHM) in ABIR. The discovery was an improvement in culture method and media for the development of *A. muciniphila* as the next generation probiotic.

## 2. Materials and Methods

### 2.1 Strains and media

#### 2.1.1 Strains

*A. muciniphila* sp. AT56 were isolated from human feces.

#### 2.1.2 Medium preparation

The BHI contained 10.0 g/L tryptone, 2.5 g/L dibasic sodium phosphate, 17.5 g/L BHI, 5.0 g/L sodium chloride, and 2.0 g/L glucose and was maintained at pH 6.5

The PM contained 4 g/L porcine mucin, 2.5 g/L disodium hydrogen phosphate, and 5.0 g/L sodium chloride and was maintained at pH 6.5(porcine mucin, Kuer, Beijing)

The HM contained 4 g/L human mucin, 2.5 g/L disodium hydrogen phosphate, and 5.0 g/L sodium chloride and was maintained at pH 6.5 (human mucin, extracted from pancreatic myxoma)

The BPM contained 10.0 g/L tryptone, 2.5 g/L disodium hydrogen phosphate, 17.5 g/L bovine heart dip powder, 5.0 g/L sodium chloride, 2.0 g/L glucose, and 4 g/L pig-derived mucin and was maintained at pH 6.5

The BHM contained 10.0 g/L tryptone, 2.5 g/L disodium hydrogen phosphate, 17.5 g/L bovine heart dip powder, 5.0 g/L sodium chloride, 2.0 g/L glucose, and 4 g/L human mucin and was maintained at pH 6.5

All media had the same concentration, magnetically stirred for 2 h, mixed well, and sterilized at 121°C for 30 min before use (Mucins remove bacteria through a filter membrane).

### 2.2. Cultivation method

#### 2.2.1 Device introduction

An *in vitro* ABIR was developed to simulate the environment of *A. muciniphila* in the real large intestine[33]. As shown in Figure 1, the bioreactor was composed of three interconnected reaction bottles with a flexible and transparent silica gel intestine that simulated the internal intestinal wall and the shape of the bionic intestine. The silica gel membrane between the reaction flask and the large intestine (Figure 1d) was filled with deionized water at 37 °C. The simulated large intestine contracted and caused peristaltic waves by on line controlling the pressure of the water flow through a circulating water pump. Thus, the materials in the simulated large intestine were mixed and moved in the system. This mixing was better than the mixing done in the fermenter, where the phase separation of fluid and solid occurred. The reaction bottle was equipped with vacuum (Figure 1g) and mixed gas (Figure 1h) control devices to continuously maintain the oxygen-free environment in the simulation chamber. The system was also added with a water absorption device that simulated the function of the large intestine (Figure 1j). The mixture in the cavity was absorbed to simulate the formation of feces. By continuously measuring the pH (Figure 1l) and secreting alkaline solution (Figure 1e) to neutralize the acid, the system pH was maintained to prevent the accumulation of microbial metabolites. The system was equipped with a dialysis device, which consisted of hollow fiber nanofiltration membranes that filtered the metabolic products of microorganisms. Thus, the microbial metabolites in the reactor cavity were maintained at physiological concentrations.

**Figure 1.**
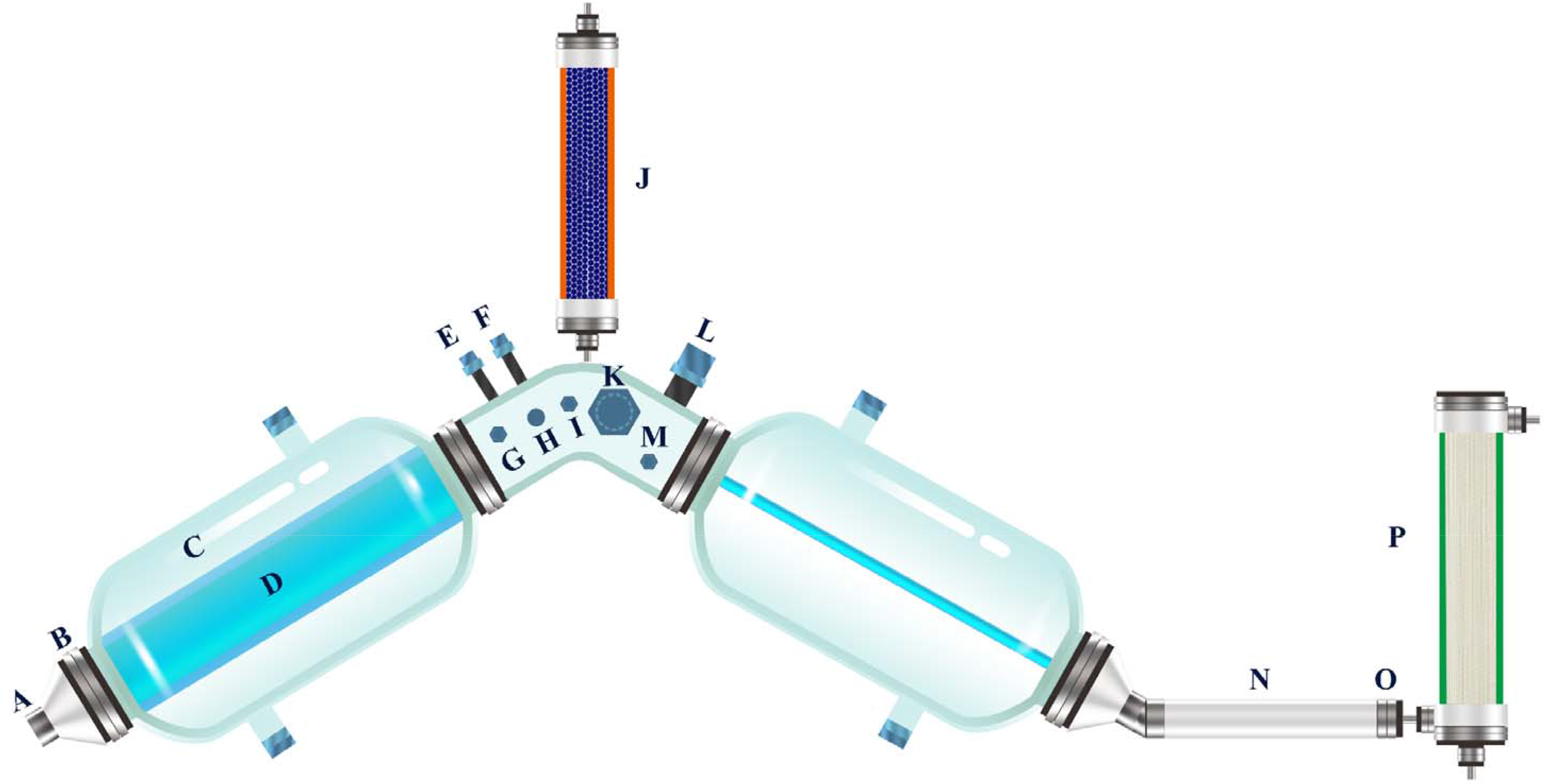
Advanced bionic intestine reactor (ABIR) : (A) discharge port, (B) sealing ring, (C) reaction bottle, (D) simulated large intestine, (E) alkali-adding port, (F) feeding port, (G) vacuum port, (H) filling mixed gas port, (I) sample port, (J) water suction device, (K) sampling port, (L) pH electrode, (M) OD_600_ detection module, (N) peristaltic tube, (O) filter screen, (P) dialysis device.

#### 2.2.2 Dynamic and static culture methods

The specific operation under dynamic culture was as follows. At first, the air tightness of the system was checked, and the whole reactor was immersed in water. The mixed gas aerator was opened to fill the reactor cavity with gas. If no bubble emerged, the air tightness of the system was good. The connection seals of each reaction bottle were detected to ensure that no bubble emerged. An aliquot of frozen stock culture of *A. muciniphila* (100 μL) was inoculated in 5 mL medium and incubated at 37 °C for 24 h under strict anaerobic conditions (hydrogen, 5%; carbon dioxide, 10%; nitrogen, 85%). The configured medium was added into the ABIR through the sample port, placed in a sterilization pot maintained at 121 °C for 30 min, and cooled to 37 °C. The extraction valve was opened, and the air in the reactor was extracted. The extraction valve was closed, and the inflation valve was opened. Anaerobic gas (hydrogen, 5%; carbon dioxide, 10%; nitrogen 85%) was poured into the reactor, and the process was repeated thrice for the reactor to have an anaerobic environment. The seed fluid flow of the incubated strain was added to the reactor cavity through a peristaltic pump. The reactor peristaltic system and the dynamic cultivation of the strain were started. In addition, the OD_600_ module (Figure 1m) can be added to this reactor to monitor the OD_600_ value of the fermentation broth in real time. For static culture, the same cultured seed liquid (5 mL) was transferred to an anaerobic bottle with BPM medium volume of 300 mL. The culture was carried out by static incubation for 48 h under the same conditions.

#### 2.2.3 Biomass determination

The fermentation broth (10 mL) was collected and centrifuged, washed twice with deionized water, moved to the centrifuge tube after weighed, and centrifuged to obtain the cells. The cells were dried at 105 °C and weighed[34].

#### 2.2.4 Determination of the short-chain fatty acids of metabolites

The fermentation broth (1 mL) was collected, added with 10 μL 2-methylbutyric acid (1 M) as an internal standard, and slowly added with 250 μL concentrated hydrochloric acid. The mixture was mixed well, added with 1 mL diethyl ether, vortexed for 1 min, and allowed to stand until the organic and the water phases separated. The supernatant (organic phase) was carefully collected, added with anhydrous sodium sulfate, vortexed, and passed through a 0.22 μm organic filter. The supernatant (5 μL) was injected into the Agilent 7890 gas chromatograph equipped with an electron capture detector (Agilent, USA), which was used to determine the short-chain fatty acids of *A. muciniphila*.

The gas chromatographic conditions were as follows: instrument, Agilent-7890A; chromatography column, HP-INNOWAX; detector temperature, 250 °C; inlet temperature, 220 °C; flow rate, 1.5 mL/min; split ratio, 1:20; heating program, 60 °C–190 °C, 4 min; and injection volume, 5 μL.

#### 2.2.5 Bacterial outer membrane protein extraction method and concentration determination

*A. muciniphila* was subjected to outer membrane protein extraction using the bacterial outer membrane protein extraction kit (BIOLABO HR0095, Beijing), and the protein content was estimated using the enhanced BCA protein assay kit (Beyotime Biotechnology, Beijing).

#### 2.2.6 FSEM and FTEM

The cultured strains were washed with PBS buffer; fixed with 2.5% glutaraldehyde solution at 4 °C for 2 h; washed again with PBS buffer; dehydrated using 30%, 50%, 70%, 80%, 90%, and 100% alcohol; dried; sprayed with gold; and lyophilized using the SU8200 (Japan) equipment. FSEM analysis was performed, and the morphology of *A. muciniphila* was observed at 3.0 KV×10.0k. The cell suspension was dropped on the copper grid and then dried at room temperature. TEM observations were taken by a transmission electronic microscope (JEM-I010, Hitachi, Tokyo, Japan) in 120 kV.

#### 2.2.7 A. muciniphila outer membrane relative thickness and diameter measurement

The TEM and SEM images were imported into the Adobe Photoshop CC 20.0.4, and the ruler tool was used to measure the relative thickness and diameter of the adventitia. Each cell was measured four times at different locations and the averaged values were presented.

#### 2.2.8 Statistical analysis

Data were expressed as mean ± SEM. Differences between the two groups were assessed using the unpaired two-tailed Student t test. The data sets that involved more than two groups were assessed using ANOVA. In the figures, data with * were significantly different at P < 0.05 in accordance with posthoc ANOVA. Data were analyzed using the GraphPad Prism version 8.00 for Windows (GraphPad Software).

Statistical comparisons were indicated with *, **, and *** for P < 0.05, P < 0.01, and P < 0.001, respectively.

## 3. Results and Discussion

### 3.1 Effects on cell growth and metabolites of A. muciniphila under static and dynamic cultures

The static culture and ABIR dynamic culture were compared in terms of biomass and metabolites. Compared with the static culture, the ABIR simulated physic-chemical and physiological processes in the real human colon, which was an idea tool for investigating the effects of the media components on *A. muciniphila* fermentation. As shown in Figure. 2a, the logarithmic growth periods of static cultures and ABIR dynamic culture were both 6–10 h, then the bacteria entered a stable stage after 12 h. The biomass under the ABIR dynamic culture increased steadily after 37 h and reached 1.92 g/L at 48 h, which was 44.36% higher than that of the static culture (1.33 g/L). In static culture, the metabolites of *A. muciniphila* accumulated partially at the late stage of the culture process and high level of SCFAs (> 80 mM) inhibited cell growth [8, 35]. In ABIR dynamic culture, simulation of real human colon peristalsis was achieved, and the cells, media and metabolites could be well mixed. Thus, the inhibition effects of metabolites could largely be released. For both static and dynamic cultures, the type and concentration of metabolites were shown in Figure. 2b. Under dynamic culture, the metabolites were mainly acetic acid, propionic acid, butyric acid, isobutyric acid, and isovaleric acid yields, which reached 47.05, 10.21, 9.87, 2.78, and 4.06 mM, respectively. The differences in yields of butyric acid between two cultures were significant (p<0.05). Simultaneously, considerable differences were observed in terms of yields of propionic acid, isobutyric acid and isovaleric acid (p<0.01). Under dynamic culture, isobutyric acid and isovaleric acid yields were relatively higher than those under static culture. In the late fermentation stage of dynamic culture, the carbon source was exhausted and *A. muciniphila* turned to protein fermentation, thereby generated corroded toxic metabolites, such as isobutyric acid and isovaleric acid[36–37].

**Figure 2.**
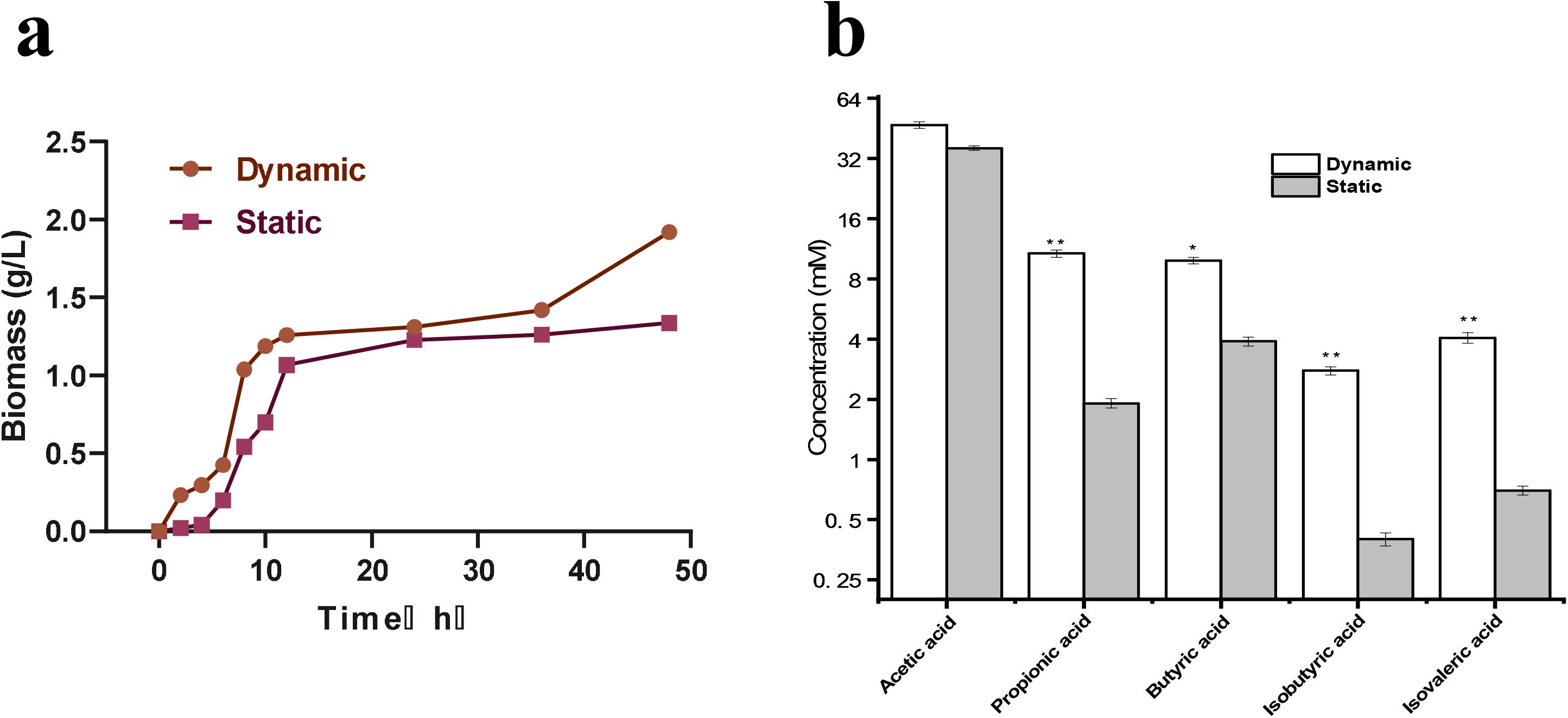
Effects on cell growth and metabolites of *A. muciniphila* under static culture and dynamic culture Note: The data are shown as the mean ± SD (n = 3) and analyzed using one-way ANOVA with Tukey’s test, *p < 0.05, **p < 0.01

### 3.2 Effects of different media on A. muciniphila growth under dynamic culture

Using five different media (BHI, PM, HM, BPM and BHM)*, A. muciniphila* biomass under dynamic cultures were compared. The mucin in the PM and HM were the only carbon and nitrogen sources for the growth of *A. muciniphila*. When *A. muciniphila* grew on these media, glycosidase was secreted to degrade the glycoprotein exposed at the terminal of mucin to produce N-acetylgalactosamine, N-acetylglucosamine, fucose, and galactose components, which could be used as nutrients for cells[38–40]. As shown in Figure 3a, using PM and HM, *A. muciniphila* grew slowly at 0–18 h, and the biomass of *A. muciniphila* grown on PM (0.68 g/L) was higher than that on HM (0.35 g/L). During this period, the secretion rates of glycosidase and protease accelerated, resulting in increased degradation of mucin. After this period, *A. muciniphila* adapted to the growth environment of medium. The growth rates of *A. muciniphila* on PM and HM were both in the logarithmic period from 18 h to 36 h. From 36–48 h, the biomass of *A. muciniphila* on HM exceeded that on PM and reached 2.89 g/L at 48 h. The results proved that *A. muciniphila* culture medium based on human mucin was an ideal medium for cell growth. Moreover, growth performances of *A. muciniphila* on BHI, BPM, and BHM were studied. As shown in Figure 3b, the *A. muciniphila* grown on BPM and BHM were in logarithmic growth phase at 4–8 h, and the *A. muciniphila* biomass on BPM (1.26 g/L) was higher than that on BHM (0.85 g/L). The *A. muciniphila* grown on BHI was in the logarithmic growth phase at 8–12 h, and *A. muciniphila* biomass of 0.61 g/L was obtained, which was lower than that on BPM and BHM. All three batches subsequently entered a stable growth phase. At 48 h, the biomass on BHI, BPM, and BHM were 1.17, 1.92, and 1.96 g/L, respectively. The BHI contained rich carbon and nitrogen sources, which supposed to fully support the growth of *A. muciniphila*. However, the biomass on BHI was the lowest among the three batches. When mucin was added in BHI, the biomass of *A. muciniphila* increased about 65%, which demonstrated that mucin was a key component for *A. muciniphila* growth. To find other cheaper sources instead of mucin, Derrien and Colleagues discovered that *A. muciniphila* could also grow on a limited amount of carbon source, including N-acetylglucosamine, N-acetylgalactosamine, and glucose[6, 31]. Based on this, a mixed medium instead of mucin to cultivate *A. muciniphila* was formed by adding glucose, soy protein, threonine, N-acetylglucosamine and N-acetylgalactosamine together. The biomass of *A. muciniphila* grown on the mixed medium reached the same level of the batches grown on BPM or BHM[14, 41]. In addition, studies found that adding fructooligosaccharides[42–43], metformin[44–45], polyphenols[46–49], probiotics[50–51] and fish oil unsaturated fatty acids[52] could increase the abundance of *in vivo A. muciniphila*.

**Figure 3.**
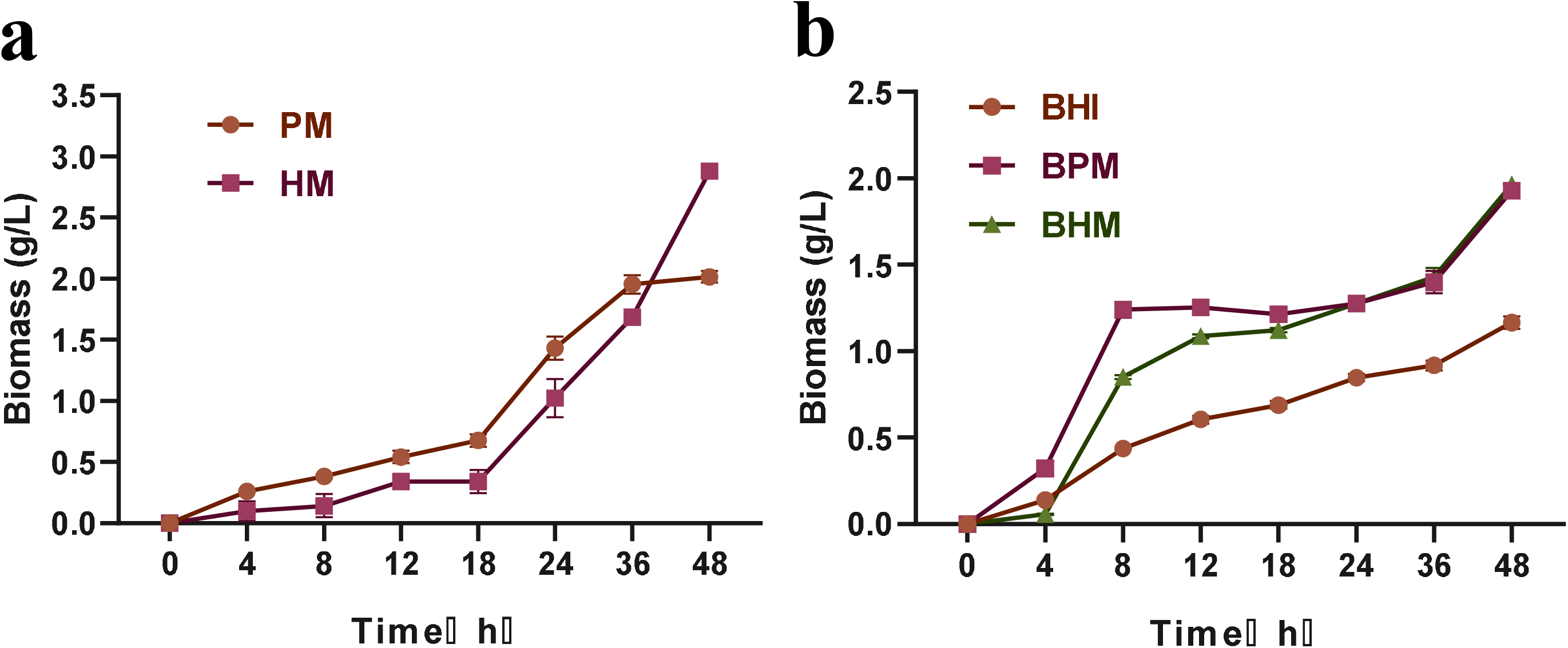
Effects of different media on *A. muciniphila* growth under dynamic culture In this figure, the brain heart infusion broth (BHI), porcine mucin (PM), human mucin (HM), BHI+ porcine mucin (BPM), and BHI+ human mucin (BHM).

### 3.3 Effects of different media on A. muciniphila metabolites under dynamic culture

Using five different media, the type and the concentration of *A. muciniphila* metabolites (SCFAs and BCFAs) were further studied, and an interesting conclusion was obtained, as shown in Figure 4. The metabolites of *A. muciniphila* on PM and HM were SCFAs (acetic acid and butyric acid), whereas the metabolites of *A. muciniphila* on BHI, BPM, and BHM were SCFAs (acetic, propionic, butyric, valeric) and BCFAs (isobutyric and isovaleric acids). For the batches grown on PM and HM, the concentrations of acetic acid (4–6 mM) were not significantly different (Figure 4b), but the concentrations of butyric acid were significantly different (Figure 4a, P < 0.05). The concentrations of butyric acid reached 1.06 mM (batch on HM) and 0.68 mM (batch on PM). For the batches grown on BHI and BHM, the concentrations of acetic acid (50.43 mM and 22.81 mM) were significantly different (P<0.05) (figure 4c). The concentrations of isobutyric acid (3.15 mM and 2.88 mM) were significantly different (figure 4e, P<0.05). The concentrations of isovaleric acid (4.25 mM and 4.07 mM) were significantly different (figure 4g, P<0.05). The concentrations of valeric acid were very significantly different (figure 4h, P<0.001). For the batches grown on BHI and BPM, the concentrations of valeric acid were very significantly different (P<0.01). For the batches grown on BPM and BHM, the concentrations of valeric acid were very significantly different (P<0.05). For the batches grown on BHI, BPM and BHM, the concentrations of propionic acid were no significant difference (figure 4d, P>0.05). The concentrations of butyric acid (5.79 mM, 9.88 mM and 12.88 mM) were significantly different (figure 4f, P<0.05). The concentrations of valeric acid (0.14 mM, 0.41 mM, and 0.51 mM) were significantly different (figure 4h, P<0.01). Studies showed that BCFAs like isobutyrate acid and isovalerate acid, were derived from the fermentation of branched-chain amino acids. In contrast to straight chain SCFAs, these compounds were considered detrimental to colonic and metabolic health [53–54]. Compared with SCFAs, these compounds were considered harmful to the colon and metabolic health. Isobutyric acid and isovaleric acid were produced on BHI, BPM and BHM. The contents of butyric acid and valeric acid on BPM and BHM were relatively high, indicating that the mucin or mucin used by *A. muciniphila* was used simultaneously with glucose. The latest research showed the causal relationship between the butyric acid produced by intestinal microorganisms and the risk of diabetes[28]. Butyric acid can promote the secretion of insulin by β cells, regulate blood sugar, and improve insulin response[36–37]. Further research was needed on the mechanism of *A. muciniphila* main metabolites (acetic acid and butyric acid) on PM and HM.

**Figure 4.**
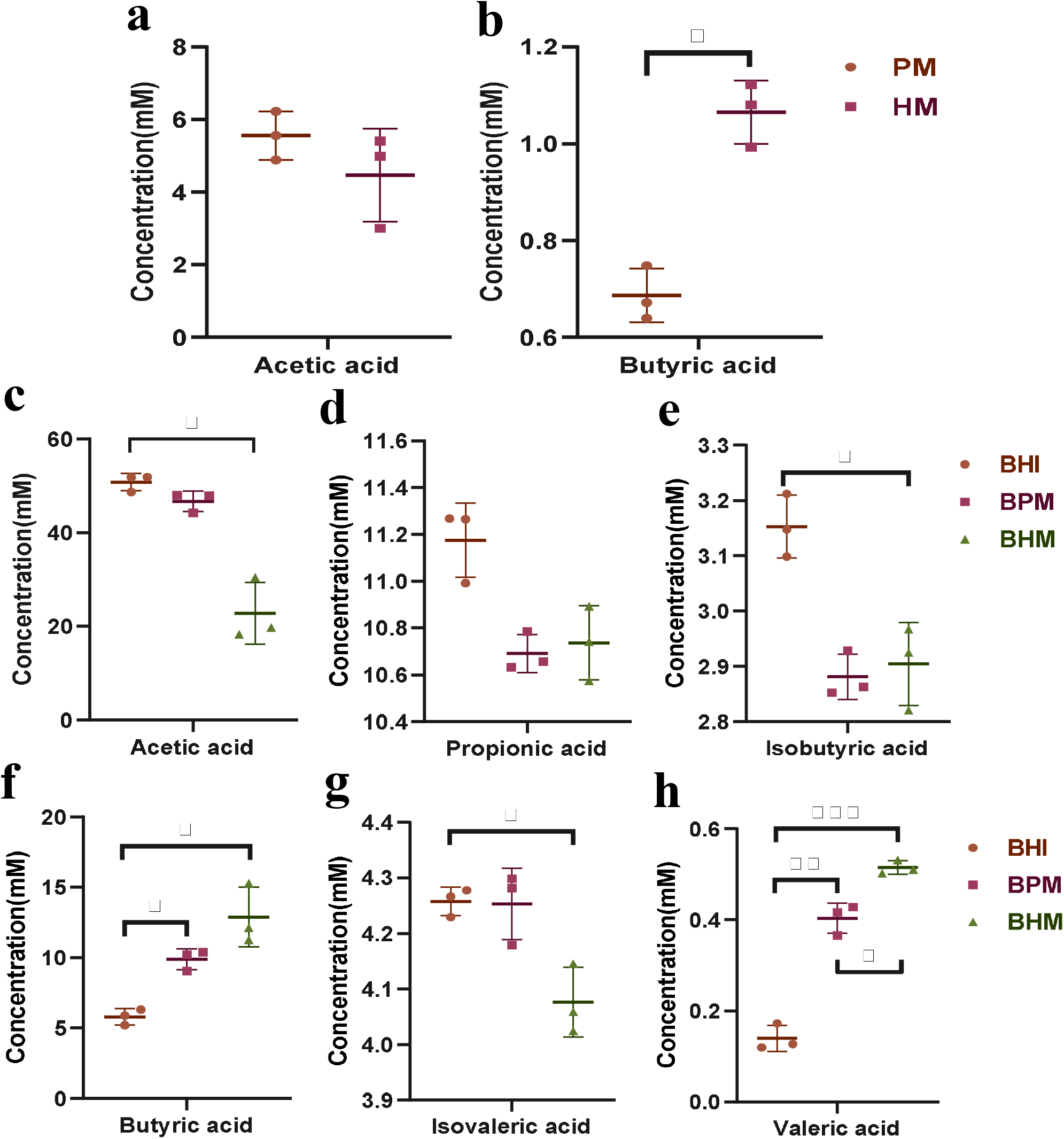
Effects of different media on metabolites of *A. muciniphila* under dynamic culture In this figure, the brain heart infusion broth (BHI), porcine mucin (PM), human mucin (HM), BHI+ porcine mucin (BPM), and BHI+ human mucin (BHM). Note: The data are shown as the mean ± SD (n = 3) and analyzed using one-way ANOVA with Tukey’s test, *p < 0.05, **p < 0.01, ***p < 0.001

### 3.4 Effects of different media on the concentration of outer membrane protein under dynamic culture

Outer membrane protein of *A. muciniphila* had a specific protein named Amuc_1100, which can maintain a stable state at the temperature of pasteurization and improve the intestinal barrier[14]. Through daily quantitative *A. muciniphila* feeding to mice, *A. muciniphila* can offset the increase in body weight and fat caused by high-fat diet (HFD) and improve glucose tolerance and insulin resistance, indicating that Amuc_1100 from pasteurization *A. muciniphila* outer membrane protein had beneficial effects on HFD-induced metabolic syndrome. Using five different media, we quantitatively detected the content of the outer membrane protein of *A. muciniphila*. As shown in Figure 5, the outer membrane protein concentrations of *A. muciniphila* in the batches grown on BHI, PM, HM, BPM, and BHM were 17.03, 24.36, 26.26, 18.34, and 19.45 μg/mg, respectively. Significant differences were observed between the batches grown on BHI and BPM (P < 0.05), BHI and BHM (P < 0.05), PM and HM (P < 0.05), PM and BHM (P < 0.05), BHI and PM (P < 0.01), HM and BHM (P < 0.01), BHI and HM (P < 0.001), PM and BPM (P < 0.001), and HM and BPM (P < 0.001). Results showed the concentration of the outer membrane protein significantly increased in the batches grown on media containing only PM or HM. For these batches, the Amuc_1100 protein of *A. muciniphila* increased accordingly. The result provided an alternative way for the improvement of Amuc_1100 protein content of *A. muciniphila* outer membrane.

**Figure 5.**
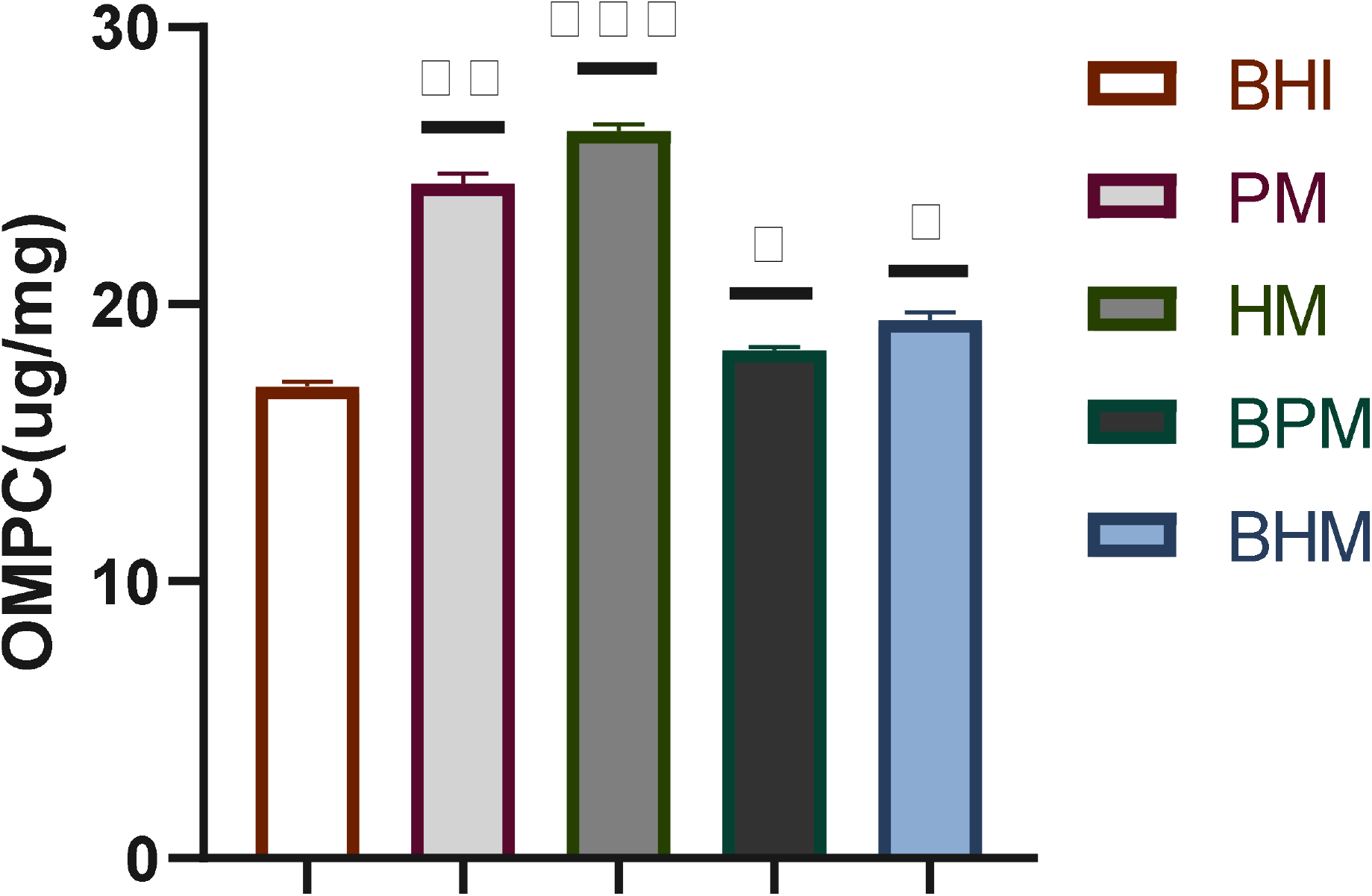
Effects of different media on concentration of *A. muciniphila* outer membrane protein under dynamic culture In this figure, the brain heart infusion broth (BHI), porcine mucin (PM), human mucin (HM), BHI+ porcine mucin (BPM), and BHI+ human mucin (BHM). Note: The data are shown as the mean ± SD (n = 3) and analyzed using one-way ANOVA with Tukey’s test, *p < 0.05, **p < 0.01, ***p < 0.001

### 3.5 Effects of different media on the thickness of A. muciniphila outer membrane and diameter of cell under dynamic culture

Figure 6 showed the TEM and SEM images of *A. muciniphila* using five different culture media under dynamic culture. The cells of *A. muciniphila* were round or elliptical. The cells of the batch grown on BHI grew alone or in pairs, whereas the cells of the batches grown on PM, HM, BPM, and BHM containing mucin grew in pairs or chains and even formed aggregates. Derrien et al. found that using porcine mucin medium under static culture[6], *A. muciniphila* could grow in single cells or in pairs and rarely grow in chains; it usually forms aggregates, and a translucent layer of material could be observed. Using BHI, this phenomenon was rarely observed, and cells appeared alone or in pairs but rarely in groups[31]. As shown in Table 1, the relative thicknesses of the outer membranes of *A. muciniphila* in the BHI, PM, HM, BPM, and BHM were 71.66, 101.20, 104.50, 71.46, and 72.11 nm, respectively. Significant differences were observed between the batches grown on BHI and PM (P < 0.05), PM and BPM (P < 0.05), PM and BHM (P < 0.05), BHI and HM (P < 0.01), HM and BPM (P < 0.01), and HM and BHM (P < 0.01). No significant difference was observed in the other remaining groups (P>0.05). The above results were consistent with those of the outer membrane protein concentration of *A. muciniphila*, indicating that increased outer membrane thickness of the cells resulted in high outer membrane protein concentration. The cell diameters of *A. muciniphila* grown on BHI, PM, HM, BPM, and BHM were 871, 985, 999, 980, and 971 nm, respectively. The cell diameter on BHI was significantly different with those on the other four media (P < 0.05), and no significant difference was observed between the other groups (P>0.05). Derrien et al. found that the cell size differed depending on the medium under static culture[6, 31]. On the mucin medium, *A. muciniphila* had a diameter and length of 640 and 690 nm, respectively. On BHI medium, *A. muciniphila* had a diameter and length of 830 nm and 1000 nm, respectively. The results were different from our discoveries because of the culture methods (the dynamic culture was carried out in this study).

**Table 1.**
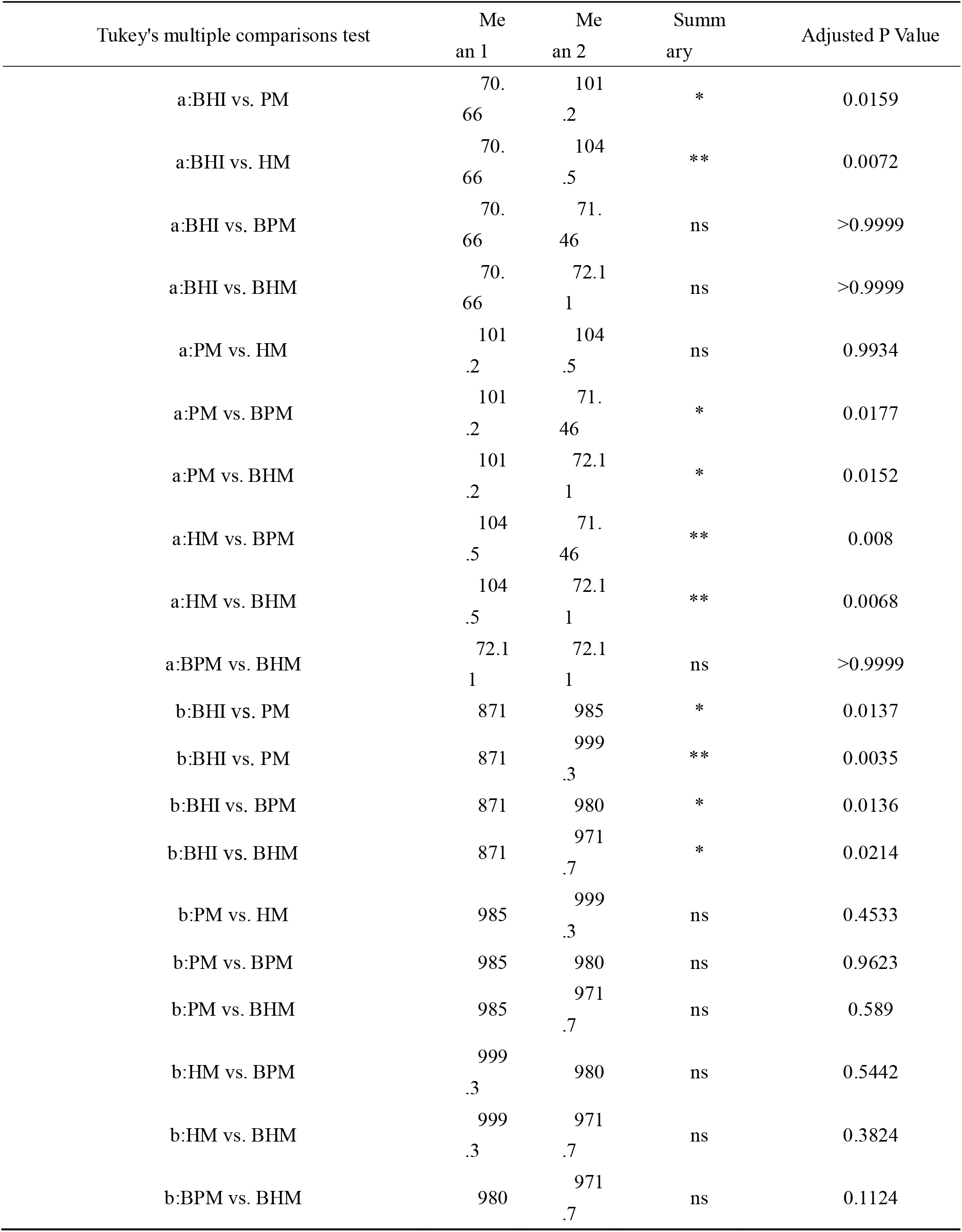
Comparative analysis of relative thickness and diameter of *A. muciniphila* outer membrane on different media under dynamic culture Note: a: Relative thickness of the outer membrane of *A. muciniphila*, b: Diameter of *A. muciniphila.* The brain heart infusion broth (BHI), porcine mucin (PM), human mucin (HM), BHI+ porcine mucin (BPM), and BHI+ human mucin (BHM). The data are shown as the mean ± SD (n = 4) and analyzed using one-way ANOVA with Tukey’s test, *p < 0.05, **p < 0.01.

**Figure 6.**
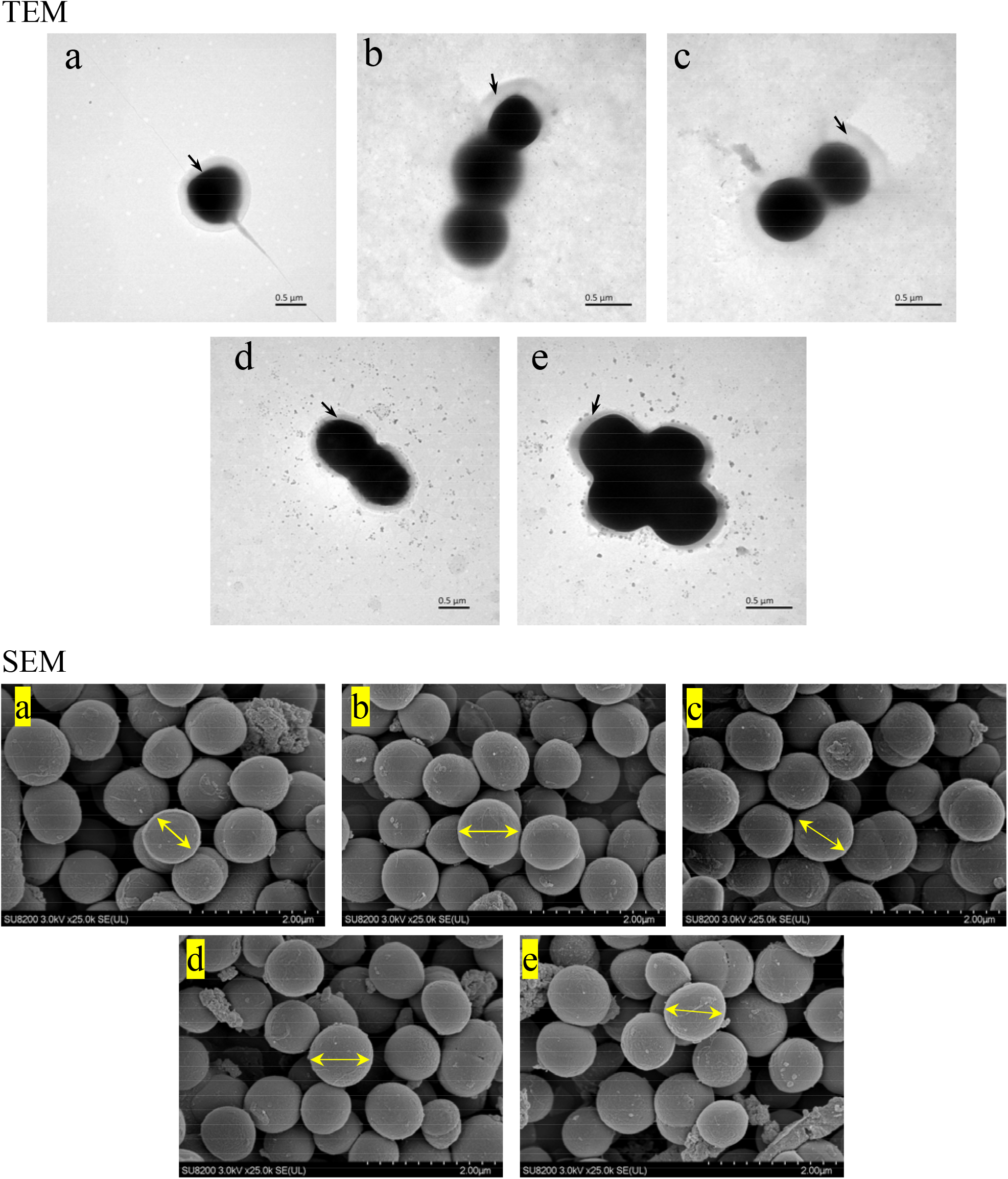
SEM and TEM images of *A. muciniphila* on different media under dynamic culture. In this figure, a, b, c, d, e represents the brain heart infusion broth (BHI), porcine mucin (PM), human mucin (HM), BHI+ porcine mucin (BPM), and BHI+ human mucin (BHM), respectively.

## 4 Conclusion

This study firstly compared *A. muciniphila* static culture in an anaerobic bottle and dynamic culture in the new biomimetic bioreactor. The *A. muciniphila* cell growth under dynamic culture improved 44.36% compared with the value under static culture using same medium. Then, the growth status, metabolites, and morphology of *A. muciniphila* under dynamic culture were further explored using five different media. The biomass of *A. muciniphila* grown on HM was the highest at 2.89 g/L, followed by that grown on PM, which indicated that mucin was an essential component for *A. muciniphila* growth. The metabolites of *A. muciniphila* in batches grown on PM and HM were acetic acid and butyric acid, and the main metabolites on BHI, BPM and BHM were acetic acid, propionic acid, isobutyric acid, butyric acid, isovaleric acid, and valeric acid. Among them, butyric acid and valeric acid on BPM and BHM were significantly different from those on BHI. The appearance of *A. muciniphila* is round or oval. The outer membrane protein concentrations of *A. muciniphila* on HM and PM were 35%–40% higher than that on the other three media. The relative thickness of *A. muciniphila* outer membrane was positively correlated with the concentration of the outer membrane protein. The diameter of *A. muciniphila* grown on BHI was the smallest (871 nm), and no significant difference in size of cells grown on the other media was observed.

## Authors’ contributions

Conceived and designed research: Min-jie Gao, Zhi-tao Li and Xiao-bei Zhan. Performed the experiments: Zhi-tao Li, Guo-ao Hu, Li Zhu and Yun Jiang. Analyzed the data: Min-jie Gao, Zhi-tao Li and Xiao-bei Zhan. Contributed reagents/materials/analytical tools: Zhi-tao Li, Zheng-long Sun, Min-jie Gao and Xiao-bei Zhan. All authors have read and approved the final manuscript for publication.

## Acknowledgements

This research was supported by National Key Research and Development Program of China (2017YFD0400302), Fundamental Research Funds for Central Universities (JUSRP51504, JUSRP51632A), and Jiangsu Province Modern Agriculture Key Project (BE2018367) and the Priority Academic Program Development of Jiangsu Higher Education Institutions, the 111 Project (No. 111-2-06) is gratefully acknowledged.

## Conflict of interest

The authors declare that the research was conducted in the absence of any commercial or financial relationships that could be construed as a potential conflict of interest.

## Notes

### Competing Interest Statement

The authors have declared no competing interest.

### Summary of Updates

The title was changed, the summary was refined, and the results and discussions were also significantly revised

